# Generation of human blastocyst-like structures from pluripotent stem cells

**DOI:** 10.1101/2021.03.09.434313

**Authors:** Yong Fan, Zhe-Ying Min, Samhan Alsolami, Zheng-Lai Ma, Ke Zhong, Wen-Di Pei, Pu-Yao Zhang, Xiang-Jin Kang, Ying-Ying Zhang, Hai-Ying Zhu, Jie Qiao, Mo Li, Yang Yu

## Abstract

Human blastocysts are comprised of the first three cell lineages of the embryo: trophectoderm, epiblast, and primitive endoderm, all of which are essential for early development and organ formation^1,2^. However, due to ethical concerns and restricted access to human blastocysts, we lack a comprehensive understanding of early human embryogenesis. To bridge this knowledge gap, we need a reliable model system that recapitulates early stages of human embryogenesis. Here we report a ∼three-dimensional (3D), two-step induction protocol for generating blastocyst-like structures (EPS-blastoids) from human extended pluripotent stem (EPS) cells. Morphological and single-cell transcriptomic analyses revealed that EPS-blastoids contain key cell lineages and are transcriptionally similar to human blastocysts. Furthermore, EPS-blastoids also exhibited the developmental potential to undergo post-implantation morphogenesis *in vitro* to form structures with a cellular composition and transcriptome signature similar to human embryos that had been cultured *in vitro* for 8 or 10 days. In conclusion, human EPS-blastoids provide a new experimental platform for studying early developmental stages of the human embryo.

**Highlights:** A method for generating human blastoids from EPS cells.

Human blastoids resemble blastocysts in terms of morphology and cell lineage composition.

Single-cell transcriptomic analyses reveal EPI, PE, and TE cell lineages in human blastoids.

Human blastoids mimic *in vitro* the morphogenetic events of pre- and early post-implantation stages.

---

Human embryogenesis begins with a fertilized egg, and then subsequent cell divisions generate the blastomere. Following activation of the embryonic genome and the beginning of compaction and polarization, the blastomere undergoes lineage segregation and morphogenetic rearrangements to form a ball-shaped structure termed the blastocyst^1,3,4^. Blastocysts contain specialized cell types, namely trophectoderm (TE), primitive endoderm (PE), and epiblast (EPI)^2,5,6^. In recent years, “multi-omic” technologies have enabled researchers to chart the gene-transcription and chromatin-modification landscapes of these cell types, providing valuable information concerning human embryogenesis^7-10^. However, the supply of human embryos is extremely limited due to ethical and technical limitations, thereby precluding a precise mechanistic understanding of early human embryogenesis. A robust *in vitro* model of human embryogenesis is urgently needed so that early human development can be systematically interrogated.

Using human embryonic stem cells (hESCs), researchers have been working toward modeling embryogenesis in a dish. Previous studies have reported that hESCs cultured in 3D soft-gel or a microfluidics device can form embryonic sac-like structures that mimic early post-implantation human epiblast and amnion development^11-13^. Recently, three-dimensional (3D) human gastrulating embryo-like structures (gastruloids) were generated by subjecting hESCs to a pulse of Wnt agonist, allowing modeling of the spatiotemporal organization of the three germ layers during

gastrulation^14^. However, all these models use only epiblast-derived hESCs and lack cells resembling the TE and PE. Therefore, they do not fully recapitulate the lineage interactions that characterize human embryonic development. Also, recent studies suggested that hESCs are of post-implantation epiblast origin and represent the primed pluripotency state^5,15^. Therefore, hESCs may not be suitable for modeling pre-implantation development.

In recent years, researchers have attempted to generate human stem cells resembling those in the pre-implantation embryo (i.e., that exhibit naïve pluripotency). These efforts revealed a continuum of pluripotency in the form of human naïve stem cells or extended pluripotency stem cells (EPS cells)^16,17^. With these cells in hand, it became possible to test whether they can self-organize into pre-implantation embryo-like structures. This interest was further stimulated by recent success in generating mouse blastocyst-like structures, termed blastoids^18-20^. Mouse EPS cell aggregates recapitulate several morphogenic hallmarks of pre-implantation embryogenesis and differentiate into both embryonic and extra-embryonic lineages, thus forming blastoids that share many features of the blastocyst. However, given the significant differences between mice and humans, it is unclear whether human EPS cells can also generate blastoids *in vitro* to mimic human embryogenesis (from pre-implantation to post-implantation stages).

In this study, we developed a three-dimensional (3D), two-step differentiation protocol for generating human blastocyst-like structures from human EPS cells (EPS-blastoids). Human EPS-blastoids resembled human blastocyst in terms of morphology, the expression of markers specific for three cell lineages, and global transcriptome signatures (obtained via single-cell RNA-sequencing). Importantly, further *in vitro* culturing of these human EPS-blastoids resulted in the emergence of structures similar to those seen in early post-implantation embryos.

## A 3D two-step differentiation method for generating human blastoid

We converted human induced pluripotent stem cells (iPSCs) into EPS cells using an established protocol^16^. We then attempted to generate human blastoids from these EPS cells using a modified version of the mouse blastoid protocol^18^. However, EPS cell aggregates treated with induction media containing BMP4 generally failed to form a cavity-containing structure after five days of induction (Extended data Fig. 1a). A few aggregates appeared to contain a small cavity, but they were enclosed by a membrane instead of TE-like cells and did not have the typical morphology and size of a blastocyst. Immunofluorescence labeling showed that the majority of these solid structures expressed the EPI marker OCT4 and the PE marker GATA6, while small fraction (∼10% of day 6 aggregates) also expressed the TE markers CK8 and GATA2/3 (about 30-50% of TE cells in each aggregate), albeit partially (Extended data Fig. 1b, c). These labeling results indicated that at least a small number of human EPS cells were capable of differentiating into the TE lineage with BMP4 induction using the modified mouse blastoid protocol^18^.

We then attempted to pretreat human EPS cells with BMP4 to generated TE-like cells. We first analyzed gene expression via real-time quantitative PCR (qPCR) during a time-course of BMP4 induction to optimize human EPS cell differentiation conditions. Between day 0 and day 5 of BMP4 stimulation, EPI-specific genes were gradually down-regulated as TE-specific genes were up-regulated (Extended data Fig. 1d). Interestingly, genes characteristic of mid- and late-TE were activated by BMP4 induction, with most TE-specific genes reaching 1.5–10-fold induction by day 3 (30- and 500-fold for *WNT7* and *GATA3*, respectively) (Extended data Fig. 1e). Higher levels of induction were observed on day 5 of BMP4 treatment. TE-like cells exhibited morphological changes by two days of BMP4 induction, with cells becoming flattened and enlarged (Extended data Fig. 1f). Immunostaining revealed more GATA2/3- and CK8-positive cells on day 3 compared with day 1 or 2 (Extended data Fig. 1g-h). As day 3 marks the onset of significant gene expression and morphological changes, we decided to use TE-like cells subjected to 3 days of BMP4 pretreatment for subsequent experiments.

In order to generate human blastoids, we first pretreated human EPS cells with BMP4 to generated TE-like cells, and then mixed human EPS cells with these TE-like cells at a ratio of 1:4∼1:5 (Fig. 1a). After 24 hours, these loosely connected cells formed small aggregates, which grew and formed structures with a small cavity by day 4. By day 5-6, blastocyst-like structures were apparent (Fig. 1h), with about 1.9% exhibiting typical blastocyst morphology (Fig. 1d). These human blastoids were morphologically similar to natural human blastocysts (Fig. 1b, c) in terms of average diameter, but blastoids seemed to have more total cells and fewer cells in the inner cell mass (ICM) (Fig. 1e-g).

**Figure 1.**
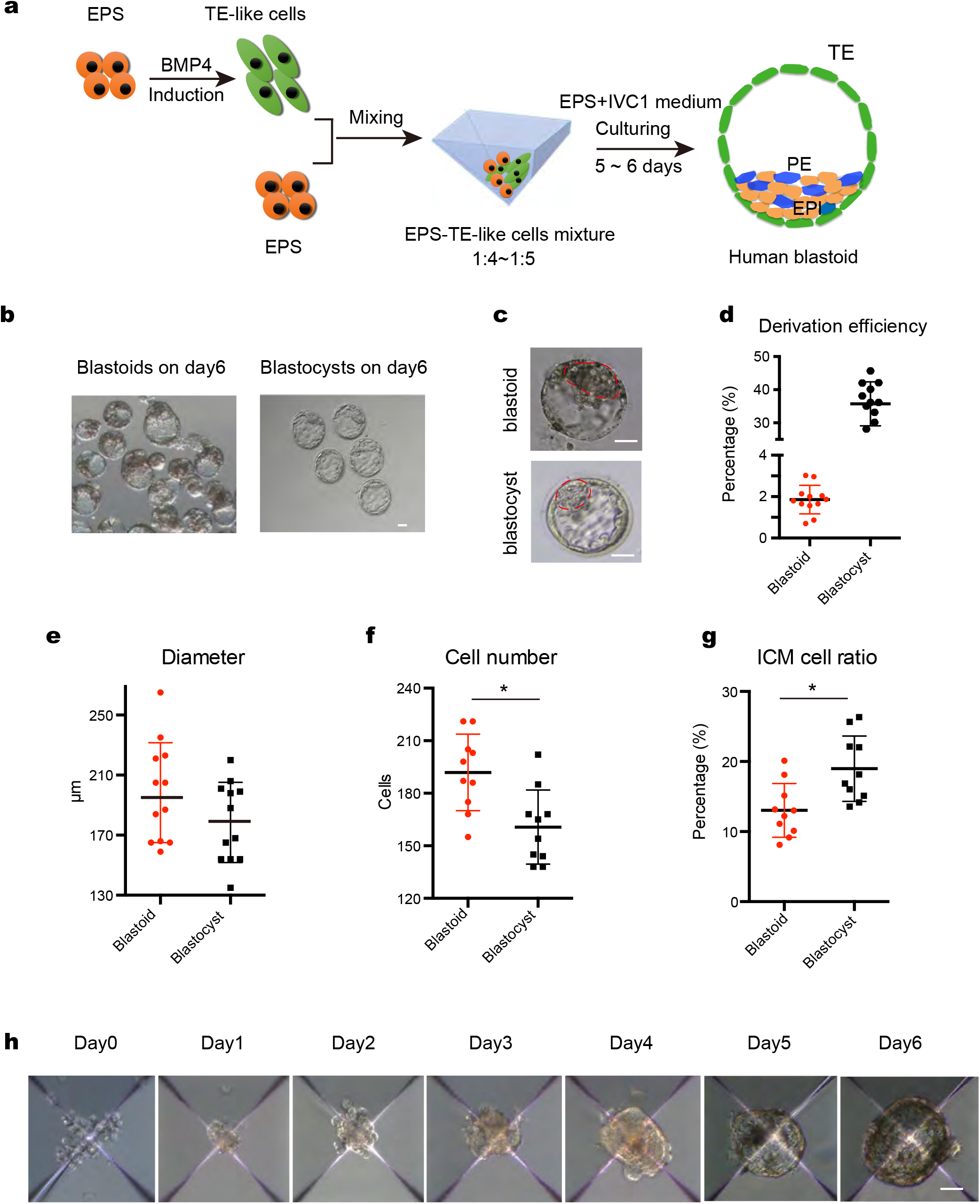
Induction of human blastoids under 3D two-step condition. a, Schematic of human blastoid formation. EPS cells were firstly induced to TE-like cells, and then TE-like cells were mixed with EPS cells and seeded together to AgreeWell on day 0. The aggregates further differentiated and organized into a human EPS blastoid. b, Phase contrast image of human blastoids on day 6, Scale bar = 50 µm. c, Phase contrast image of human EPS-blastoid (upper) and human blastocyst (lower). Red line indicated inner cell mass (ICM) of the structure and the outer layer cells represented trophoblast cells (TE). Scale bar = 50 µm. d, Derivation efficiency of human blastoids is about 1% that significantly lower than the developmental efficiency of human blastocysts. e-g, Mean diameter (e), total cell number (f) and ICM cell ratio (g) were quantified between human EPS-blastoids and blastocyst. n= 30 EPS-blastoids, n= 30 blastocyst. h, Phase-contrast images of human aggregates in the indicated days showing the formation of human blastoids from day 0 to day 6. Scale bar = 50 µm. Data in c, data are mean ± SD (n = 12 blastoids). \*\**P*< 0.001. Data in e, data are mean ± SD (n = 12 blastoids). *P*> 0.05. Data in f and g, data are mean ± SD (n = 12 blastoids). \**P*< 0.05. Data in h, data are numbers (n = 40 blastoids, 40 blastocysts), \*\**P*< 0.001.

## Human EPS blastoids contain cells of the three blastocyst lineages

Mixing TE-like cells and EPS cells resulted in cell aggregation and the formation of blastocyst-like structures (EPS-blastoids) on day 5. To investigate whether EPS-blastoid formation recapitulated key cellular processes, we monitored cellular dynamics of blastoids during day 2 and 3. We found that the cell-adhesion protein, E-cadherin (E-cad), localized to cell-cell junctions, indicating cell-cell interactions and communication within EPS aggregates^21^.

We sought to determine whether blastoids contained the three cell lineages of the blastocyst, namely EPI, PE, and TE, as all three are necessary for an embryo to develop beyond implantation. Immunofluorescence analysis of day 4 blastoids revealed extremely low levels of OCT4 in the outer cell layer (TE cells), with highest levels localized to EPI cells within the interior of the ICM. Cells surrounding these OCT4-positive cells expressed the PE-marker, GATA6, as seen in natural early blastocysts^22^ (Extended data Fig. 2a). Immunofluorescence analysis of day 5 and 6 EPS-blastoids revealed that OCT4-positive cells localized exclusively to the ICM-like compartment (Fig. 2a, f), whereas cells in the outer layer expressed the TE-specific transcription factors, GATA2 and GATA3 (Fig. 2d,). The outer layer of cells also expressed the trophoblast-specific cytokeratin, KRT8 (CK8), indicative of TE specification (Fig. 2e, f). The PE-lineage marker, GATA6, localized to cells adjacent to the OCT4-positive cells (not those within the outer layer of the blastoid) (Fig. 2c, g). This pattern of localization is similar to that seen in human blastocysts (Fig. 2h).

**Figure 2.**
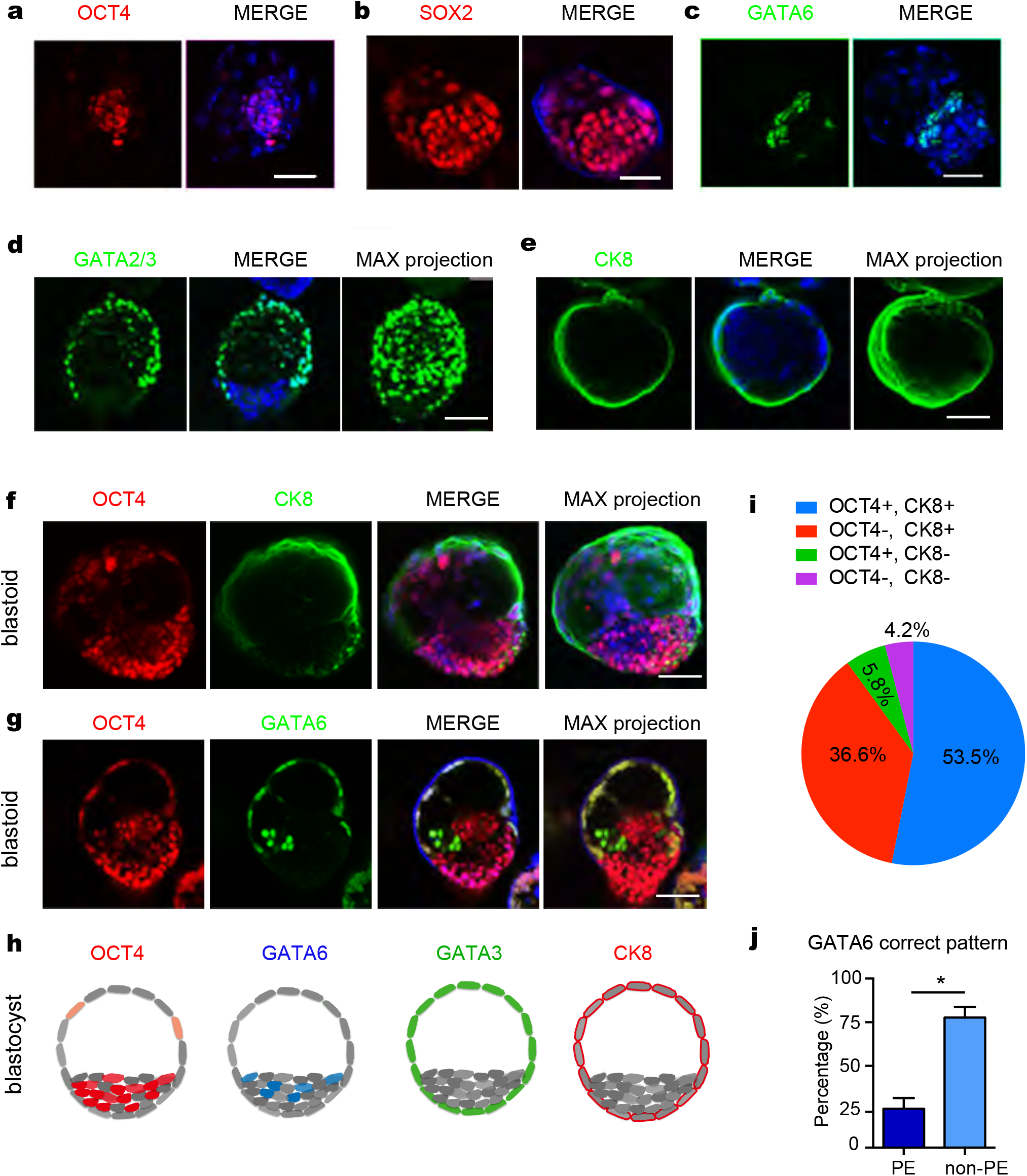
Expression of specific development-related markers. a-g, Immunofluorescence staining of human EPS-blastoids for EPI lineage marker (OCT4, SOX2), TE lineage markers (CK8 and GATA2/3), and PE lineage marker (GATA6), Scale bar = 50µm. h, Model of immunofluorescence staining of human blastocyst for OCT4 in EPI, GATA6 in PE, GATA2/3 and CK8 in TE lineages, Scale bar = 50 µm. i, j, Quantification of the percentage of human blastoids with correct allocation of OCT4 and CK8 (i) and GATA6 (j). Data in i, data are mean ± SD (n = 172 blastoids). \**P*< 0.05. Data in j, data are mean ± SD (n = 131 blastoids).

We then calculated the percentage of CK8-positive or OCT4-positive cells in day 6 EPS-blastoids (n = 172). These analyses revealed a smaller fraction of OCT4-positive cells than seen in human blastocysts, whereas the fraction of CK8-positive cells was reminiscent of human blastocysts. On average, there were approximately 15% and 80% of cells expressed OCT4 and CK8 in one blastoid, respectively (Extended Data Fig. 2a-b). Of these, 53.5% exhibited the correct pattern of ICM-(OCT4+) and TE-like (CK8+) localization, 36.6% exhibited only correct TE-like pattern (CK8+), 5.8% had only the ICM-like lineage, and 4.2% exhibited mislocalization of ICM- and TE-like pattern (Fig. 2i). With further development of early blastocysts, ICM divided two lineages: EPI and PE. We examine whether blastoid could develop into a late blastcyst-like structure composed three lineages (EPI, PE, and TE). Of 131 blastoids on day 6, about 8% cells were shown GATA6 positive expression in one blastoid on average (Extended Data Fig. 2c). Moreover, the ratio of blastoids that expressed OCT4, CK8 or GATA6 were all around 80% (Extended Data Fig. 2d-f), and 26% showed PE-like lineage (GATA6+, in a stochastic manner) (Fig. 2m). Using this induction system, a large proportion of EPS cells failed to form blastoids, although they expressed markers indicative of the EPI, PE, and TE lineages. They instead retained features of day 4 aggregates (Extended Data Fig. 3a). In some abnormal day 6 aggregates, OCT4, SOX2, or GATA6 localized improperly to TE cells (Extended Data Fig. 3b-e) instead of the ICM. In summary, these results demonstrated that EPS-blastoids recapitulated the segregation of TE and ICM cell lineages, and that these blastoid structures possessed the three lineages typical of a blastocyst.

To confirm the existence of EPI and TE lineages in these EPS-blastoids, we attempted to derive ESCs and trophoblast stem cells (TSCs) from day 6 EPS blastoids. We were able to establish 2 ES cell lines from 5 EPS-blastoids, and 3 TS cell lines from 10 EPS-blastoids using the culture condition reported previously^23,24^. ESCs and TSCs were morphologically similar to those derived from blastocyst (Extended Data Fig. 4a and f). ESC colonies expressed the pluripotency markers OCT4, SOX2, SSEA4, and TRA-1-60 (Extended Data Fig. 4b-e), whereas TSC colonies expressed the TE-specific markers GATA3, CK7, and TFAP2C (Extended Data Fig. 4g-l).

## Single-cell transcriptome analysis of human blastoids

We performed single-cell RNA-sequencing (scRNA-seq) analysis of 200 day 6 human blastoids. After quality control and filtering, 10,000 single cells were further analyzed using the bioinformatic suite Seurat. Uniform manifold approximation and projection (UMAP) clustering analysis showed that cells from human blastoids could be divided into 20 clusters. Base on the expression of marker genes (Extended Data Fig. 5a), we characterized seven clusters as ICM/EPI, three cluster as PE, and three clusters as TE (Fig. 3a). The remaining seven clusters seemed to express both ICM and TE markers and could each represent an intermediate cell type (Fig. 3a). In addition, we performed unsupervised clustering analysis and confirmed that the top genes specific for each lineage were consistent with the UMAP clustering results. The ICM/EPI cluster expressed *OCT4* (*POU5F1*) and *NANOG*, the PE cluster expressed *GATA6* and *PDFGRA*, and the TE cluster expressed *TFAP2C* and *GATA2* (Fig. 3b). We then performed differential gene expression analysis to identify gene-expression differences between the lineages. A total of 822 genes were enriched in the ICM/EPI lineage. The PE lineage was characterized by 680 genes, and 1264 genes were differentially expressed in the TE lineage (Fig. 3c-e),.

**Figure 3.**
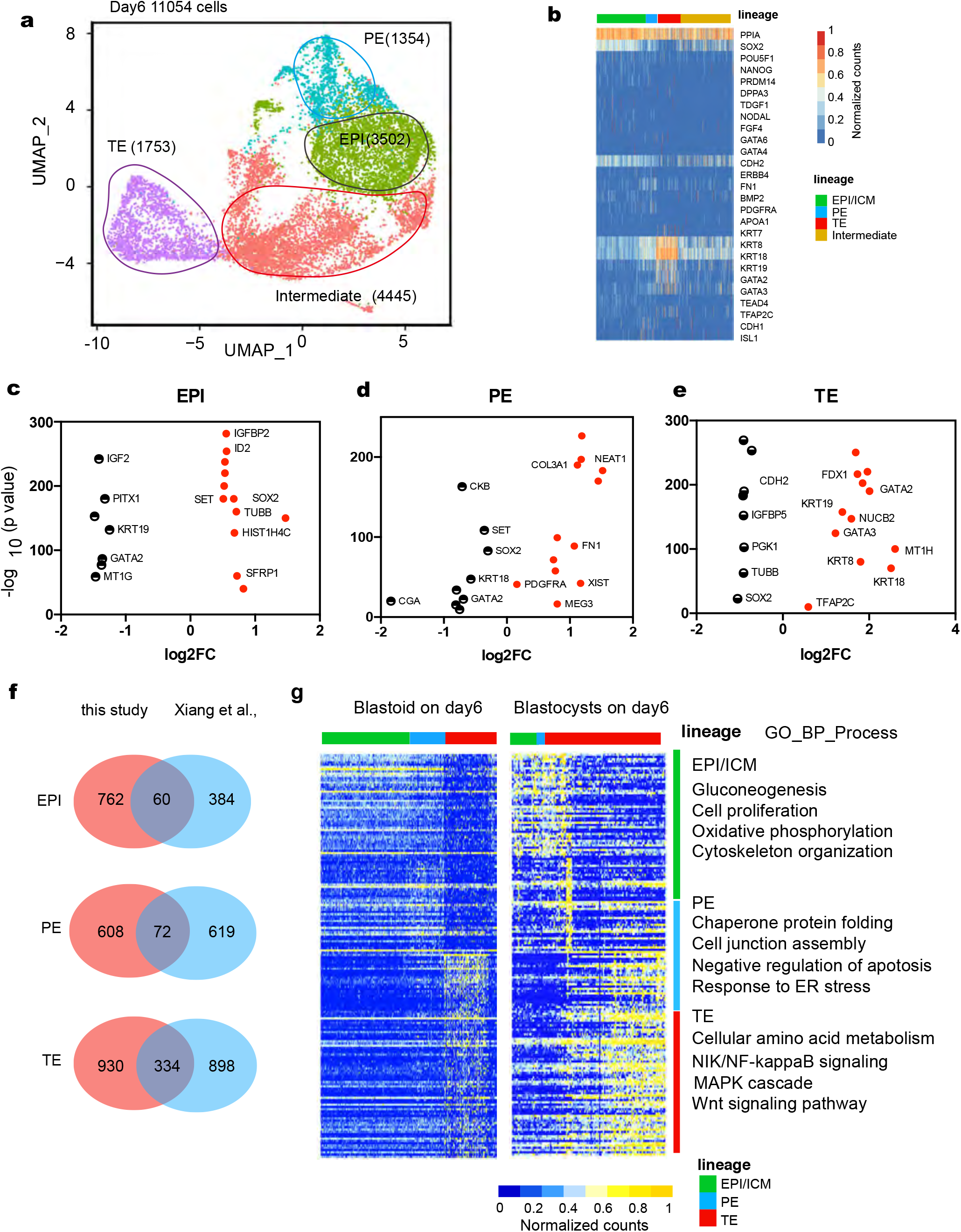
Landscape of transcriptome in human blastoids on day 6. a, A UMAP plot 2867 cells from human blastoids showing the cells in blastoids were divided into 4 major clusters on day 6. PE, ICM/EPI, TE, and intermediate subgroups were determined according to the lineage specific markers. b, Heatmap of lineage signature genes expression on day 6. c-e, Dot plots showing the top differentially expressed genes specifically in EPI, PE or TE lineage. f, g, Comparison of overlapping lineage specific genes between our results and previous study (Xiang et, al) (f). Overlapping genes expression between our results and previous study was also performed by heatmap and GO term analysis (g).

To reveal similarities and differences between EPS-blastoids and natural blastocysts, we compared our scRNA-seq data from day 6 EPS-blastoids with two independent datasets acquired from human blastocysts^7,25^. Comparisons with Xiang’s data derived from embryos 6–7 days post fertilization (d.p.f.) revealed 60, 72, and 334 genes that overlapped with those in the EPS-blastoid EPI, PE, and TE clusters, respectively (Fig. 3f, Supplementary Table1). Hierarchical clustering showed comparable expression pattern across the three lineages of day 6 blastoids and the 6–7 d.p.f. embryos^25^ (Xiang et al.)(Fig. 3g). Comparisons of our results with data derived from 5–7 d.p.f. embryos^7^ (Petropoulos et al.) revealed 64, 27, and 77 genes that overlapped with those in EPS-blastoid EPI, PE, and TE clusters, respectively (Extended Data Fig. 5b, Supplementary Table2). We also performed gene ontology (GO) enrichment analysis for the two sets of overlapping genes by lineages. GO terms were largely similar between the two analyses (Extended Data Fig. 5c). Overall, our scRNA-seq analysis revealed comparable transcriptome landscapes for EPS-blastoids and early blastocysts, and confirmed that by day 6 human blastoids contained the three cell lineages found in blastocysts.

## Human EPS blastoids can develop post-implantation embryonic structures

To test whether human EPS-blastoid could undergo post-implantation morphogenesis, we kept culturing day 6 blastoids within 2 and 4 days (hereafter referred day 8 and day 10 embryonic structures) using a previously established *in vitro* culture (IVC) system, which needed matrigel-coated plate and modified IVC1/2 medium to mimic blastocyst implantation^26,27^. On day 8, GATA6-positive cells encircled the OCT4-positive cells (Fig. 4a). On day 10, the localization patterns for OCT4 and GATA6 resembled day 8, except for an increased number of cells within the day 10 embryonic structures (Fig. 4b).

**Figure 4.**
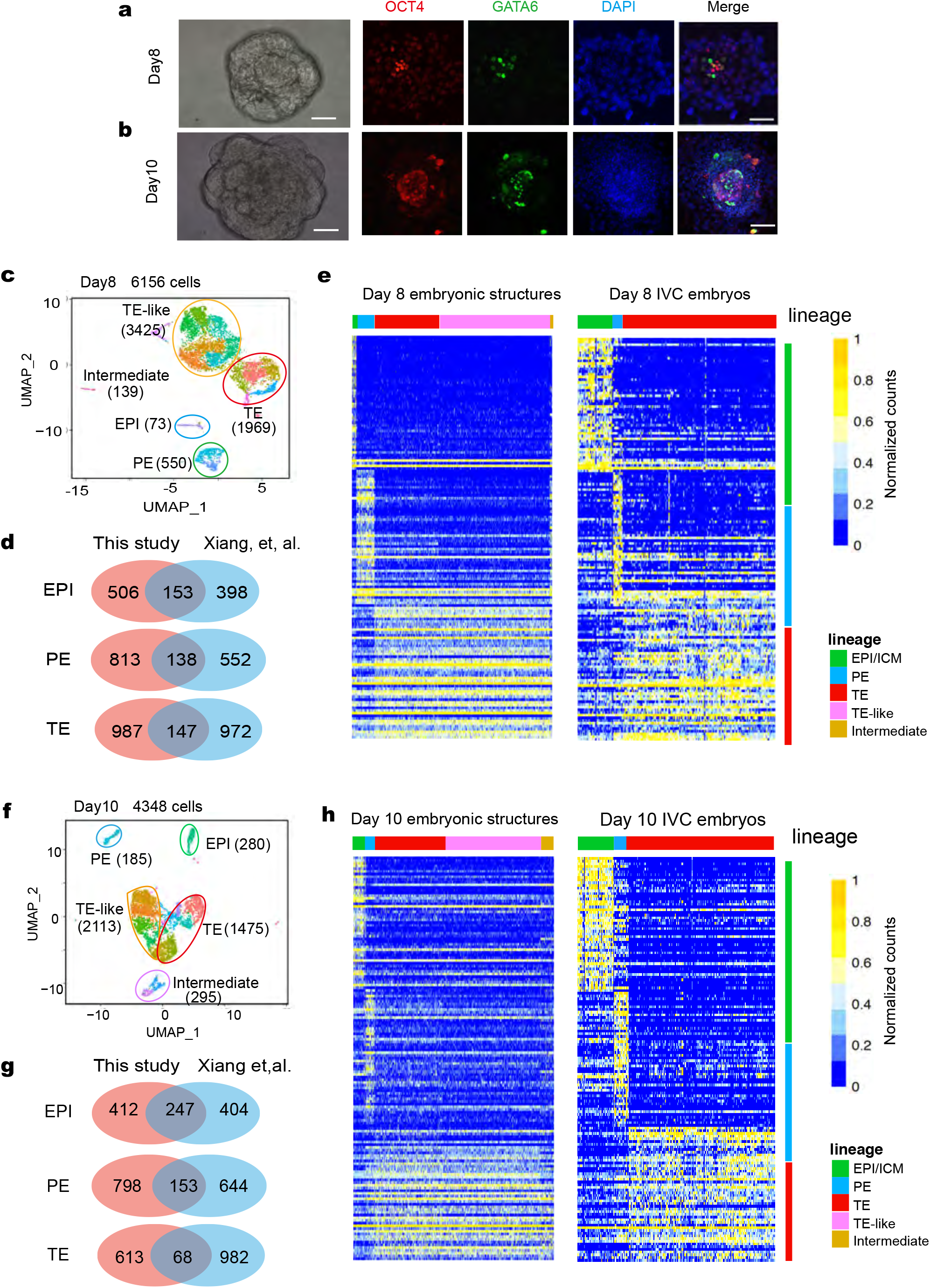
Landscape of transcriptome in human embryo-like structures from blastoids on day 8 and day 10. a, b, Immunofluorescence staining of EPI marker OCT4 and PE marker GATA6 in Day 8 (a) and Day 10 (b) embryonic structures. c, A UMAP plot analysis revealed 5 clusters in embryonic structures on day 8, identified as ICM/EPI, PE, TE, TE-like and intermediate cluster. d, Comparison of overlapped lineage specific genes between our results of embryonic structures on day 8 and previous studies (data of 7–9 d.p.f. in Xiang et al.). e, Heatmap of lineage specific genes overlapping between embryonic structures on day 8 and blastocysts IVC embryos (Xiang et al. data of 7–9 d.p.f. embryos). f, A UMAP plot analyses revealed 5 clusters in embryonic structures on day 10, identified as ICM/EPI, PE, TE, TE-like and intermediate cluster. g, Comparison of overlapping lineage specific genes between our results of embryonic structures and previous studies (Xiang et al. data of 9–12 d.p.f. embryos). h, Heatmap of lineage specific genes overlapped between embryonic structures on day 10 and blastocysts IVC embryos (Xiang et al. data of 9–12 d.p.f. embryos).

We then performed scRNA-seq analysis using day 8 (40) and day 10 (20) human embryonic structures. After quality control and filtering, 6156 single cells from day 8 and 4348 single cells from day 10 were further analyzed. UMAP analysis by Seurat revealed that the primary clusters could be identified as ICM/EPI, PE, TE, TE-like, and intermediate on day 8 (Fig. 4c, Extended Data Fig. 6c) and day 10 (Fig. 4f, Extended Data Fig. 7c). This is based on the expression of 12 representative marker genes (Extended Data Fig. 6a, 7a). We further confirmed these clustering result with an independent lineage score analysis, which showed the formation of three major clusters on day 8 and 10 (Extended Data Fig. 6b, 7b). The human placenta consists of three major trophoblast (TE) subpopulations: cytotrophoblasts (CTBs), extravillous cytotrophoblasts (EVTs), and syncytiotrophoblasts (STBs). Accordingly, we were able to subdivide the TE subpopulation from day 8 EPS embryonic structures into pre-CTBs, early EVTs, and early STBs clusters (Extended Data Fig. 6d, e). For day 10 EPS embryonic structures, we identified an additional subgroup of cells representing EVTs (Extended Data Fig. 7d, e).

To reveal similarities and differences between EPS embryonic structures and natural blastocysts subjected to IVC, we compared our results of EPS embryonic structures from day 8 and 10 with data acquired from 8–12 d.p.f IVC embryos^25^ (Xiang et al.). A total of 153, 138, and 147 genes, specific for our ICM, PE, and TE clusters (day 8) overlapped with Xiang’ s analyses of 7–9 d.p.f IVC embryos (Fig. 4d). Similarly, comparison of day 10 embryonic structures and 9–12 d.p.f. IVC embryos revealed 247, 153, and 68 genes that overlapped with our EPI, PE and TE clusters (Fig. 4g), respectively. Hierarchical clustering and GO analysis showed that our EPS embryonic structures (day 8 and 10) and 8–12 d.p.f IVC embryos had a similar transcriptional profile (Fig. 4e, h). In conclusion, embryo-like structures cultured from EPS-blastoids *in vitro* (day 8 or 10) resembled natural human embryos in terms of their single-cell transcriptome landscape, confirming their potential for modeling early post-implantation development.

## Discussion

Mouse EPS cells can be induced to form blastocyst-like structures (blastoids)^18,19^, but human blastoids have not yet been reported. Because of the significant differences between mouse and human developmental processes, it is thought that the generation of human blastoids may be more challenging. Indeed, applying the modified mouse culture system to human EPS cells failed to generated blastoids. To overcome this obstacle, we developed a 3D, two-step induction system for generating blastoids from human EPS cells. In our two-step induction system, we first exposed EPS cells to BMP4 for 3 days to induce TE-like cells formation. These TE-like cells were then mixed with EPS cells to generate EPS-blastoids. We found that TE-like cells expressed early-, mid-, and late-TE cell markers, which enhanced their subsequent developmental potential. Human EPS-blastoids were similar to human blastocysts of the same stage based on both morphology and cell lineage analysis. The latter conclusion was based on immunofluorescence and scRNA-seq analyses. The differentiation efficiency of EPS-blastoids was lower than seen for mouse blastoids (1.9% vs. 15% in the human and mouse systems, respectively). One potential reason for this difference is the difficulty in maintaining stemness and pluripotency of human EPS cells in our differentiation system. The human EPS cells were difficult to keep dome-shaped clones during the cultivation, which contained differentiation cells disturbing blastoids formation. Other reasons include differences between human and mouse early embryonic development, and differences between the mouse and human embryo culture system. Culture medium is an important component for inducing blastoids. The efficiency of the culture system for human embryos is not as robust as the mouse system. Up to 90% of mouse embryos can develop to blastocysts *in vitro*, whereas only 50% of human embryos reach the blastocyst stage. Also, the comparisons of our scRNA-seq data with previous data suggested that the human two-step blastoid differentiation system must be further optimized to achieve an efficient and reliable model system for studying the human blastocyst.

Our human EPS-blastoids recapitulated to a great extent the 3D-architecture of human fertilized blastocysts, and exhibited all three developmental lineages. Functionally, we could derive both ESCs and TSCs from human EPS-blastoids. More importantly, they gave rise to post-implantation structures. These observations suggested that hESP-blastoids manifest at least some functionalities of the natural human blastocyst. The ability of hEPS-blastoid to generate several types of mature trophoblasts that are similar to the human placenta, offers great promise for studying placenta disorders in the future.

In summary, we have established for the first time an *in vitro* system for generating human blastoids that accurately recapitulate the development of a human blastocyst. Human blastoids provide an alternative and potentially high-throughput platform for exploring the mechanisms of human blastocyst development and stem cell differentiation during pre- and post-implantation stages.

## Acknowledgements

The work was supported by the National Key R&D Program of China (2016YFC1000601, 2019YFA0110804, 2018YFA0801400, 2018YFC1003203), the National Natural Science Funds (81971381, 81771580, 81571400, 82071723, 81871162), and Outstanding Overseas Returnees Fund of the Peking University Third Hospital (No. BYSYLXHG2019002).

## Author contributions

YF and ZYM performed majority of the experiments related to blastoid derivation and identification. ZYM, EZ, SA and ZLM modified the blastoid derivation system. XJK and HYZ performed the human blastocyst collection. ML, ZYM, WDP and PYZ performed the bioinformatics analysis. EZ and ZLM performed the blastoids *in-vitro* culture experiments. ML, YY, TT, YF and ZYM analyzed the data and wrote the manuscript. YY, TT, ML and YF conceived and supervised the study.

## Methods

### Human samples and ethics statement

Human skin fibroblasts were isolated from chest of a female aborted fetus that were obtained with informed written consent and approval by the Third Affiliated Hospital of Guangzhou Medical University. The generation of induced pluripotent stem cells with donated human fibroblasts was approved by the Ethics of Third Affiliated Hospital of Guangzhou Medical University. Human blastocysts produced from in vitro fertilization for clinical purposes were got with informed written consent and approval by the Third Affiliated Hospital of Guangzhou Medical University. All procedures were approved by the Institutional Review Board of the Third Affiliated Hospital of Guangzhou Medical University (2020027), and Peking University Third Hospital (S2020022).

### Generation of human EPS cells from iPSCs

Human iPSCs were generated using the electroporation (4D-Nucleofector System, Lonza) of fibroblasts with episomal vectors, including pCXLE-hOCT3/4-shp53-F, pCXLE-hSK and pCXLE-hUL as reported previously^28^. The derived iPSCs were cultured on mitomycin C-treated MEF feeder cells in human ESC medium. The human ESC medium consisted of DMEM/F12 (Thermo Fisher Scientific, 11330-032) supplemented with 20% Knockout Serum Replacement (Thermo Fisher Scientific, A3181502), 0.1 mM non-essential amino acids (Thermo Fisher Scientific, 10828-028), 2 mmol/L GlutaMAX (Thermo Fisher Scientific, 35050-061), 0.1 mmol/L β-mercaptoethanol (Thermo Fisher Scientific, 21985-023), 1% penicillin/streptomycin (Thermo Fisher Scientific, 21985-023), and 10 ng/mL recombinant bFGF (Thermo Fisher Scientific, PHG0261). Human iPSCs were converted to EPS cells referred to protocol described previously^16^. Human iPSCs were digested into single cells by TrypLE Express Enzyme (Thermo Fisher Scientific, 12604021) or Accutase (Stem Cell Technologies, #07920) and seeded to the plate with ICR MEF feeders. After 12 hours seeding, the human ESC medium was replaced with human N2B27-LCDM medium. N2B27 basal medium was prepared as following: 240 mL DMEM/F12 (Thermo Fisher Scientific, 11330-032), 240 mL Neurobasal (Thermo Fisher Scientific, 21103-049), 2.5 mL N2 supplement (Thermo Fisher Scientific, 17502-048), 5 mL B27 supplement (Thermo Fisher Scientific, 12587-010), 1% nonessential amino acids (Thermo Fisher Scientific, 11140-050), 1% GlutaMAX (Thermo Fisher Scientific, 35050-061), 0.1 mM β-mercaptoethanol (Thermo Fisher Scientific, 21985-023), 1% penicillin-streptomycin (Thermo Fisher Scientific, 15140-122), 5% knockout serum replacement (KSR, Thermo Fisher Scientific, A3181502). Human N2B27-LCDM medium consisted of N2B27 basal medium supplemented with 10 ng/mL human LIF (R&D Systems, 7734), 3µM CHIR99021 (Tocris, 4423), 2µM (S)-(+)-Dimethindene maleate (Tocris, 1425), and 2µM minocycline hydrochloride (Selleck, S4226), 1 µM IWR endo-1 (Selleck, S7086), and 2 µM Y-27632 (Selleck, S1049). The N2B27-LCDM medium was changed every day. Dome-shaped colonies emerged after 3-6 days. Then the cells were dissociated and passaged to the next generation.

### In vitro 3D generation of human EPS-blastoids

The cells culturing conditions were as follows: 37°C, 5% CO2 and saturated humidity. Human EPS cells were differentiated with BMP4 for 3-4 days. On day 0, the cultured EPS cells were digested into single cells by TrypLE Express, and the feeder was removed by pasting twice on 0.5% gelatin for 15-20 minutes each time. The EPS cells were collected, centrifuged and seeded into plate pre-treated with Matrigel in BMP4 differentiation medium. After 3 days of differentiation, cells were dissociated into single cells with 0.05% trypsin. Mix the BMP4 treated cells (1.0 × 10^5^ cells) and EPS cells (2.0 × 10^4^ cells) together following 5:1 ratio to total 1.2 × 10^5^ cells per well with 0.5 ml culture medium, and then seeded into one well of 24-well aggreWell400 culture plate pretreated with anti-adherence rinsing solution following instruction (Stem Cell Technologies, #07010). The culture medium was slightly changed without disturbing aggregates in the V-shape bottom every day and aggregates were collected on day 6. The culture medium of human EPS-blastoids is composed of EPS medium and IVC1 medium in a ratio of 1.5:1 (v/v). BMP4 differentiation medium was composed of N2-B27 basal medium, supplemented with 25ng/ml BMP4 (R&D SYSTEMS, 314-BP-010), 2µM Y-27632 (Selleck, S1049). IVC1 culture medium is composed of Advanced DMEM/F12, 20% Heat-inactivated FBS, 2mM L-glutaMAX (Thermo Fisher Scientific, 35050-061), 100U/mL penicillin and streptomycin (Thermo Fisher Scientific, 15140-122), 1% ITS-X (Thermo Fisher Scientific, 51500-056), 1% sodium pyruvate , 8nM β-estradiol (Sigma-Aldrich, E8875), 200ng/mL progesterone (Sigma-Aldrich, P0130) and 25 µM N-acetyl-L-cysteine (Sigma-Aldrich, A7250).

### Derivation of ESCs and TSCs from human blastoids

To derive ES cells^23^, individual EPS-blastoid was transferred onto a MEF layer in a 96-well plate and cultured with human ESC culture medium (see above). After 2-3 days, EPS-blastoid attached to the MEF and outgrew. Then outgrowth was digested by TrypLE Express and transferred into a new MEF feeder layer. Colony was picked, digested, and seeded onto a new MEF plate for ESC deriveation. To derive TS cells^24^, EPS-blastoid was seeded in a 96-well plate coated with 5 µg/mL Collagen IV (Coring, 354233) at 37°C overnight and cultured in human TS medium (DMEM/F12 supplemented with β-mercaptoethanol, 0.2% FBS, 0.5% penicillin-streptomycin, 0.3% BSA, 1% ITS-X supplement, 1.5 µg/mL L-ascorbic acid (Wako, 013-12061), 50 ng/mL EGF (Wako, 053-07871), 2 µM CHIR99021, 0.5 µM A83-01, 1 µM SB431542 (Wako, 031-24291), 0.8 mM VPA (Wako, 227-01071) and 5µM Y27632). When cells reached 70% confluence, cells were digested by TrypLE Express and transferred into a new Collagen IV-coated 96-well plate at a ratio of 1:4.

### In vitro culture of EPS-blastoids for 8 and 10 days

The method of embryo extended cultured *in vitro* refers to the previous studies ^26,27^. The blastoids were collected on day 6, and transferred to the 8-well plate (treated with Matrigel 30 minutes in advance), and cultured with IVC1 for 2 days. On day8, observe whether blastoid was adherent to the bottle of the dish or not, and replace the culture medium with IVC2 for further culture if the blastoid was adherent to the wall till to day 10. The composition of IVC1 medium is the same as describe in the “In vitro 3D generation of human EPS-blastoids”, IVC2 culture medium is composed of Advanced DMEM/F12, 30% Knockout serum, 2mM L-glutamine, 100U/mL penicillin and streptomycin, 1% ITS-X, 1% sodium pyruvate,8nM β-estradiol, 200ng/mL progesterone, 2 µM Y27632 and 25 µM N-Acetyl-L-cysteine.

### Immunofluorescence labeling

The samples were fixed with 4% paraformaldehyde in phosphate buffered saline (PBS) for 20 min at room temperature, washed three times with PBS, and permeabilized with 0.2% Triton X-100 in PBS for 15 min. After blocking with 5% BSA in PBS for 2h at room temperature, samples were then incubated with primary antibody diluted in blocking buffer overnight at 4°C. After primary antibody incubation, samples were washed three times with PBS containing 0.1% Tween20. Samples were washed three times with PBS containing 0.1% Tween20 and incubated with fluorescence-conjugated secondary antibodies diluted in blocking buffer at temperature for 2h. Nuclei were stained with Hoechst 33342 (Sigma, 94403) at 1µg/mL. Zeiss LSM 710 or 880 confocal microscope were used for imaging. Images were processed by ZEN (Zeiss) and Fiji (ImageJ, V2.0.0) softwares. The primary antibodies and dilutions were following: mouse anti-OCT4 (Santa Cruz, sc5279, polyclonal, A0616, F2719, 1:200), rabbit anti-GATA6 (Cell Signaling Technology, 5851S, monoclonal, D61E4, 4, 1:1000), mouse anti-SOX2 (Abcam, ab171380, monoclonal, 20G5, GR32833, 1:200), rabbit anti-GATA2/3 (Abcam, ab182747, monoclonal, EPR17874, GR22065, 1:200), rabbit anti-CK8 (Abcam, ab53280, monoclonal, EP1628Y, 2, 1:200), mouse anti-E-cadherin (Abcam, ab1416, monoclonal, HECD-1, GR33004, 1:200), rabbit anti-CK7 (Abcam, ab192077, monoclonal, EPR1619Y, GR325605, 1:200). The secondary antibodies were: Alexa Fluor 488 Goat anti-Rabbit IgG (H+L) (Thermo Fisher Scientific, A-11008), Alexa Fluor 555 Goat anti-Mouse IgG (H+L) (Cell Signaling Technology, 4409S), Alexa Fluor 647 Goat anti-Rabbit IgG (H+L) (Abcam, ab150083, GR3269213).

### Real-time quantitative PCR

Total RNA was extracted using TRIzol (Invitrogen, 15596018). RNA (2 µg) was reverse-transcribed to cDNA template using RevertAid First Strand cDNA Synthesis Kit (Thermo Fisher Scientific, K1621). Q-PCR was analyzed in the Applied Biosystems QuantStudio 3. The changes of genes were calculated by the comparative ΔΔCt method. The primer sequences were shown in supplemental table1. All the experiments were performed in triplicates.

### Single-cell RNA-sequencing

Human EPS-blastoids were picked up by mouth pipette and washed with PBS containing 0.05% BSA. About day 6 EPS-blastoids (200), day 8 (40) and day 10 (20) human embryonic structures were collected for single-cell RNA-seq. Samples on day 6 were dissociated with enzyme mix composed of 0.5×Versene (Lonza, 17711E), 0.5×Accumax (STEMCELL Technologies, 07921) and 0.05×DNaseI (STEMCELL Technologies, 07900) at 37°C for 30 min with agitation and terminated by 5% BSA in PBS. Samples on day 8 or 10 were dissociated with enzyme mix composed of 0.25% Trypsin (Thermo Fisher Scientific, 25200056), and 0.05×DNaseI (STEMCELL Technologies, 07900) at 37°C for 15 min with agitation and terminated by 5% BSA in PBS. Dissociated cells were repeated pipetting and washed with PBS containing 0.05% BSA. Using single cell 3 ’Library and Gel Bead Kit V3 (10×Genomics, 1000075), the cell suspension (300-600 living cells per microliter determined by Count Star, about 20,000 dissociated cells each sample) were loaded onto the Chromium Single Cell B Chip (10×Genomics, 1000074) and processed in the Chromium single cell controller (10×Genomics) to generate single-cell gel beads in the emulsion according to the manufacturer’ s protocol. In short, single cells were suspended in PBS containing 0.04% BSA. No more than 10,000 cells were added to each channel, and the target cell recovery rate was estimated to 8,000∼9,000 cells. Captured cells were lysed and the released RNA was barcoded through reverse transcription in individual GEMs (Zheng et al., 2017). Using a S1000TM Touch Thermal Cycler (Bio Rad) to reverse transcribe, the GEMs were programed at 53°C for 45 min, followed 85°C for 5 min, and hold at 4°C. The cDNA was generated and then amplified, and the quality was assessed using the Agilent 4200. According to the manufacture’ s instruction, Single-cell RNA-seq libraries were constructed using Single Cell 3’ Library and Gel Bead Kit V3. Finally, sequencing was performed on the Illumina Novaseq6000 sequencer with a sequencing depth of at least 60,000 reads per cell and 150 bp (PE150) paired-end reads (performed by CapitalBio, Beijing).

### Analysis of single-cell RNA-sequencing data

The Cell Ranger software was obtained from 10×Genomics website https://support.10×genomics.com/single-cell-gene-expression/software/downloads/latest. Alignment, filtering, barcode counting, and UMI counting were performed with cellranger count module to generate feature-barcode matrix and determine clusters. Dimensionality reduction was performed using PCA and the first ten principle components were used to generate clusters by K-means algorithm and graph-based algorithm, respectively. The other clustering method is Seurat 3.0.12 (R package). The R package Seurat 3.0.12 was used to analyze feature-barcode matrix as following steps: 1, Cells whose gene numbers were less than 200, or unique features counts over 9000, or gene numbers ranked in the top 1%, or mitochondrial gene ratio was more than 15% according to quality control matrix plots, were regarded as abnormal and filtered out. 2, UMI counts were normalized with NormalizeData function by default settings. A non-linear dimensionality reduction was performed using PCA, clustered with resolution setting at 1-2, and visualized by TSNE and UMAP. 3, Differentially expressed genes (DEGs) in clusters were determined by the FindAllMarkers function. For the DEGs, the GO terms in biological process were enriched using DAVID. Heatmap of DEGs between clusters were performed with R package pheatmap.

### Statistical analysis

Statistical analyses were performed with GraphPad Prism 8 software, using unpaired two-tailed Student’ s t-tests and one-way ANOVA. All of the statistical tests performed are indicated in the figure legends. The data are presented as mean ± SD, and P<0.05 was regarded as significant differences. For cell numbers and gene expression, the significant differences between two samples were analyzed by GraphPad Prism 8 software.

### Data availability

Single-cell RNA-seq data have been deposited in the Gene Expression Omnibus (GEO) under accession number GSE158971 (scRNA-seq data website: https://www.ncbi.nlm.nih.gov/geo/query/acc.cgi?acc=GSE158971) the single-cell RNA-seq of human pre- and early post-implantation embryos (for Figs. 3-4) are with GEO accession GSE136447 (ref.^25^) and E-MTAB-3929 (ref.^7^).

