## Supplemental figures for "Generation of human blastocyst-like structures from pluripotent stem cells"

**Extended Data Figure 1 BMP4 induced the specific markers of TE lineage**

a, Upper panel: Schematic of aggregate formation in mouse blastoid system (without TE-like cells mixing). Lower panel: aggregate morphology. b, Immunostaining of aggregates generated in mouse system for EPI marker OCT4 and PE marker GATA6. Scale bar = 50 μm. c, Immunostaining of aggregates generated in mouse system for TE marker CK8 and GATA2/3. Scale bar = 50 μm. d, Gene expressions of three lineage markers under BMP4 treatment from Day 0 to Day 5 analyzed by qPCR. e, Quantification of lineage specific genes from day0 to day5 under BMP4 treatment. f, Morphology of TE-like cells in indicated induction days (day 1 to day 3). g-h, Immunostaining of EPI marker (OCT4 and SOX2), PE marker (GATA6) and TE marker (GATA2/3 and CK8) under BMP4 treatment from Day1 to Day3. Scale bar = 50 μm.

**Extended Data Figure 2 Analysis of specific lineage markers expression**

a-c, The proportion of OCT4 (a), GATA6 (b), and CK8 (c) positive cells in human blastoids and blastocysts. d-f, Quantification of the percentage of human EPS aggregates with OCT4 (d), GATA6 (e) and CK8 (f) expression. g, Immunostaining of E-cadherin expression in human aggregates on day 2 (up panel) and day 3 (down panel). Data in a and b, data are mean ± SD (n = 172 blastoids). $^*P< 0.05$. Data in c, d, e and f, data are mean ± SD (n = 131 blastoids). $^*P< 0.05$.

**Extended Data Figure 3 Summary of lineage specific marker expression in human blastoids under different induction system**

a, Immunostaining of human blastoid on day 4 for EPI marker OCT4 and PE marker GATA6. OCT4 was expressed in the center of human blastoids, GATA6 was expressed surround the OCT4 positive cells. b-e, The blastoids with the abnormal expression or allocation of the lineage markers OCT4, SOX2, GATA2/3 and GATA6.

**Extended Data Figure 4 Derivation of ESCs and TSCs from human EPS- blastoids**

a, Phase contrast image of human ESCs derived from human EPS-blastoid. b-e, Immunostaining of human ESCs derived from human EPS-blastoid for pluripotency marker OCT4 (b), SOX2 (c), SSEA4 (d) and TRA-1-60 (e). f, Phase contrast image of human TSCs derived from human EPS-blastoid. g-i, Immunostaining of human TSCs derived from human EPS-blastoid for trophoblast related marker GATA3 (g), TFAP2C (h) and CK8 (i).

**Extended Data Figure 5 Lineage-specific gene expression characteristics in blastoids on day 6 by single-cell RNA-seq**

a, The expression of 12 representative lineage-related genes of blastoids on Day 6 shown in UMAP plot, including ICM/EPI, PE and TE lineage. b, Comparison of overlapped lineage specific genes between our results and previous study (Petropoulos et al.). c, Overlapped genes in three lineages were annotated by GO Biological_Process enrichment.

**Extended Data Figure 6 Lineage-specific gene expression characteristics in human embryonic structures on day 8**

a, The expression of 12 representative lineage-specific genes of embryonic structures on day 8 shown in UMAP plot, including EPI, PE and TE. b, Lineage identification of embryonic structures on day 8 was confirmed by lineage score analysis. The lineage scores for each cell were performed by ggtern plot, calculated using lineage specific genes. The individual cells are colored by the lineage identities. c, Heatmap of lineage signature gene expression on day 8, such as OCT4/NANOG for ICM/EPI, GATA6 for PE, and GATA3 for TE. d, A UMAP plot analysis revealed 3 sub-clusters in TE cluster (including early-EVTs, early-STBs and post-CTBs) of day 8 embryonic structures. e, Heatmap of top specific expressed genes in TE clusters on day 8.

**Extended Data Figure 7 Lineage-specific gene expression characteristics in human embryonic structures on day 10**

a, The expression of 12 representative lineage-specific genes of embryonic structures on day 10 shown in UMAP plot, including EPI, PE and TE. b, Lineage identification of human embryonic structures on day 10 was confirmed by lineage score analysis. c, Heatmap of lineage signature genes expression on day 10. d, A UMAP plot analysis revealed 4 sub-clusters in TE cluster (including early-EVTs, early-STBs, post-CTBs and EVTs) of embryonic structures on day 10. Compared with day 8, EVTs cluster can be identified in embryonic structures on day 10. e, Heatmap of top specific expressed genes in TE clusters on day 10.

**Supplementary Table1. Summary of GO Term analysis of overlapped genes for each lineage between EPS-blastoids on day 6 and blastocysts on day 6–7 (Xiang et al.)**

**Supplementary Table2. Summary of GO Term analysis of overlapped genes for each lineage between EPS-blastoids on day 6 and blastocysts on day 5–7 (Petropoulos et al)**

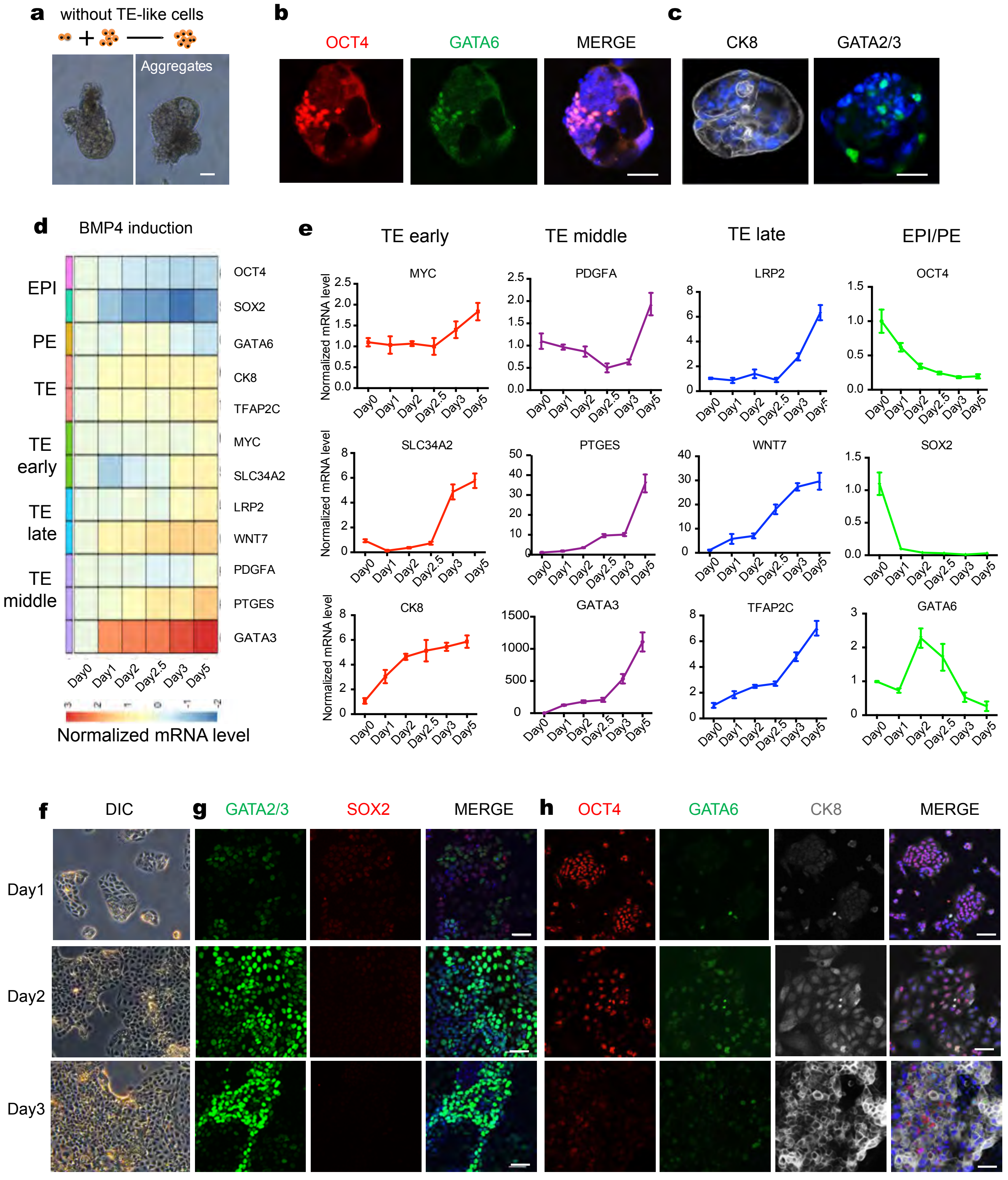


Extended Fig 1

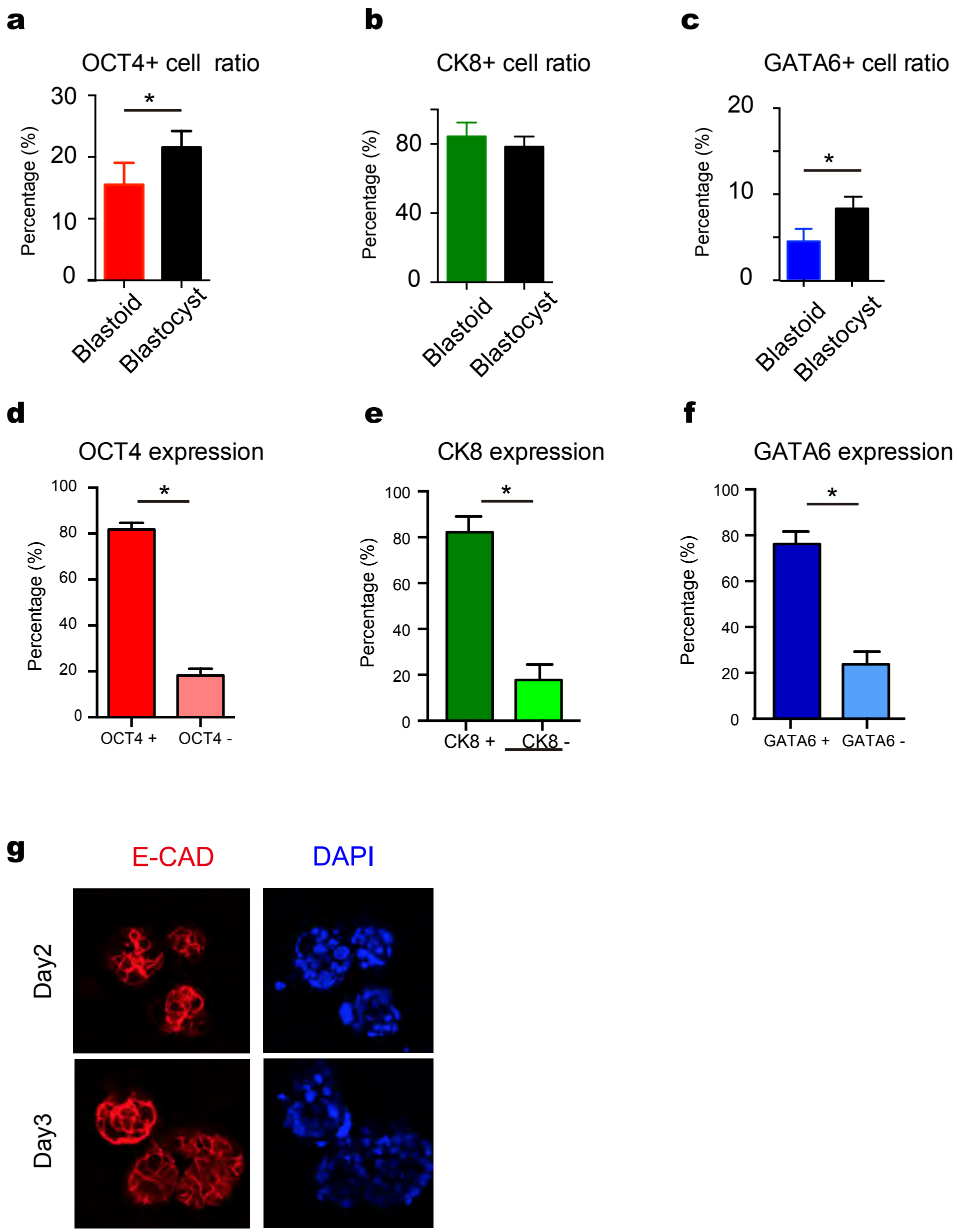


Extended Fig 2

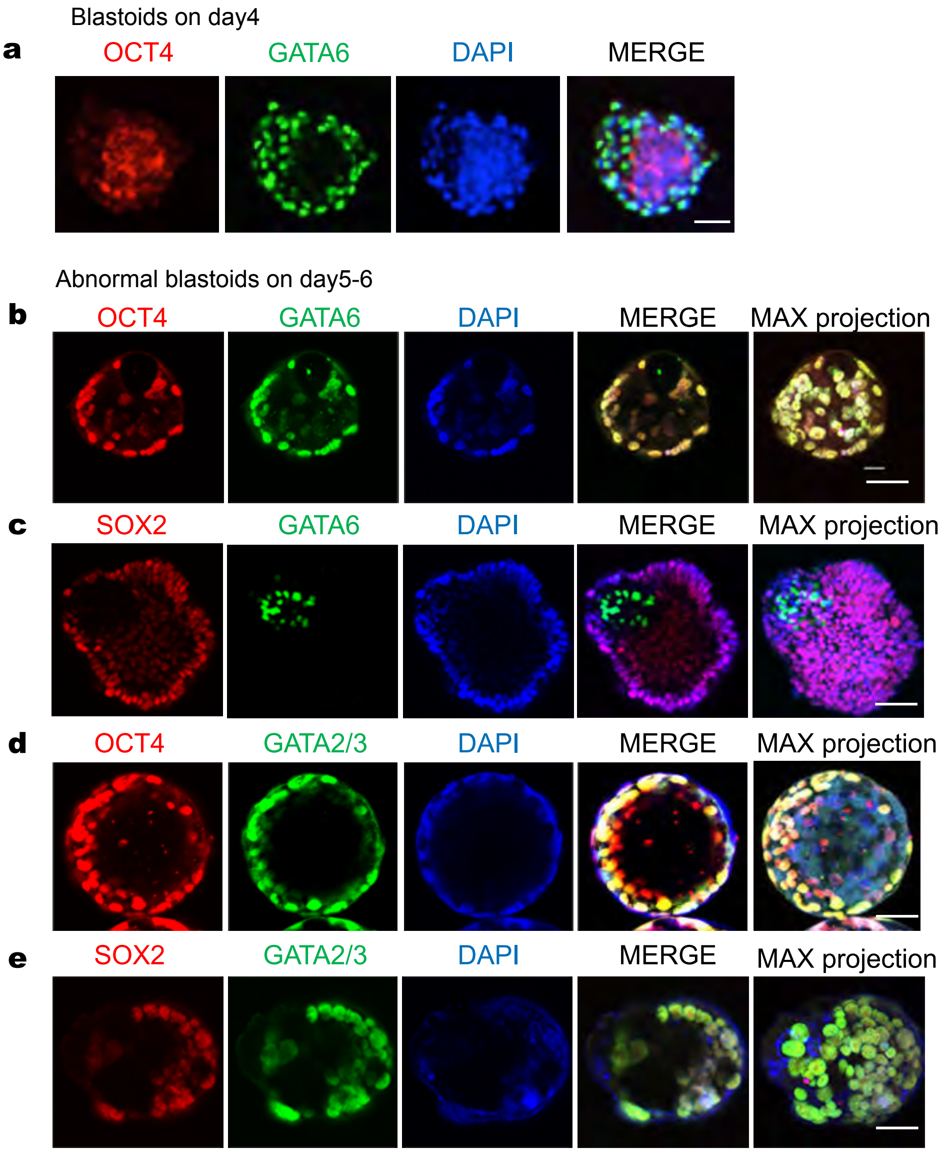


Extended Fig 3

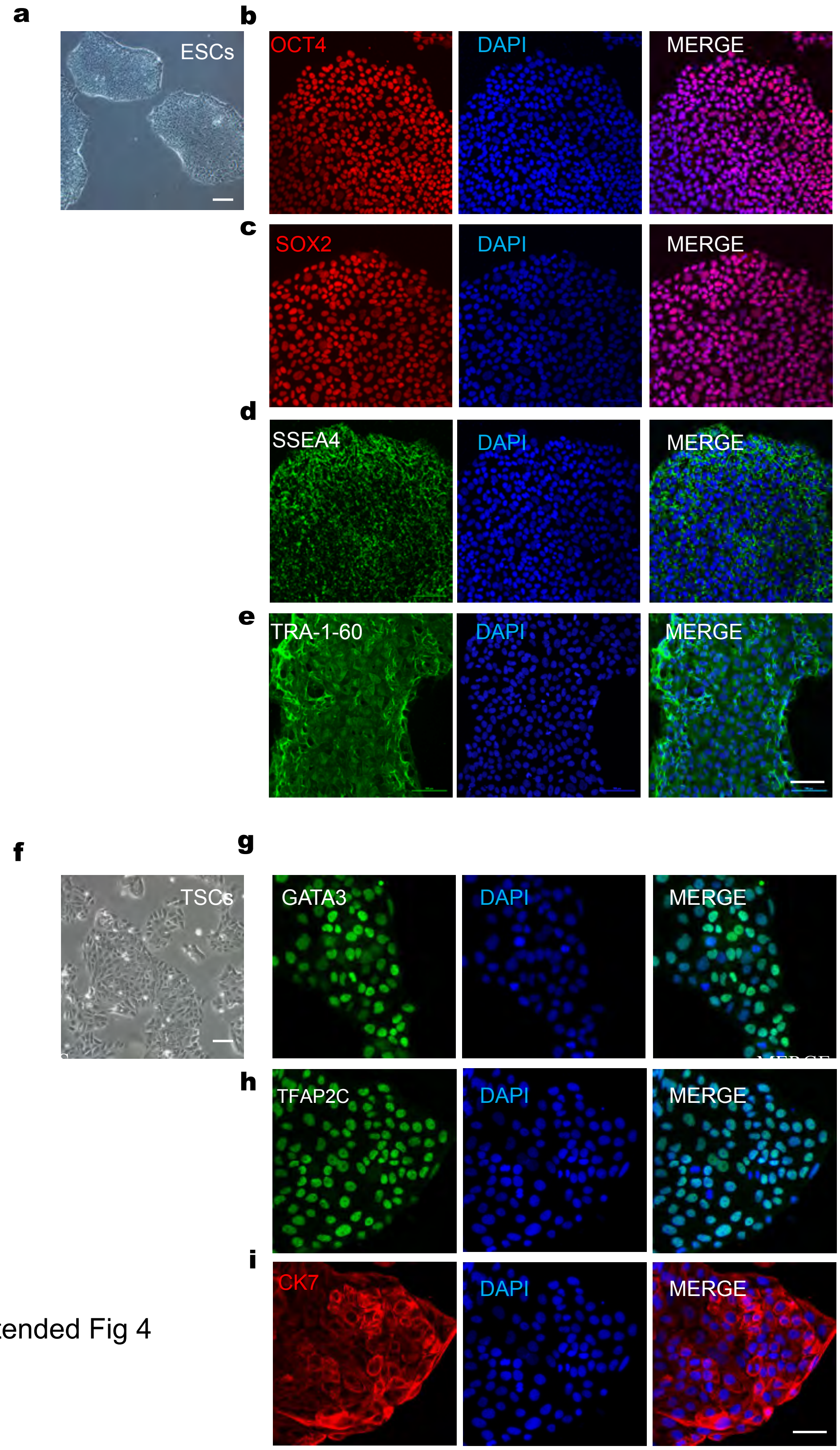


Extended Fig 4

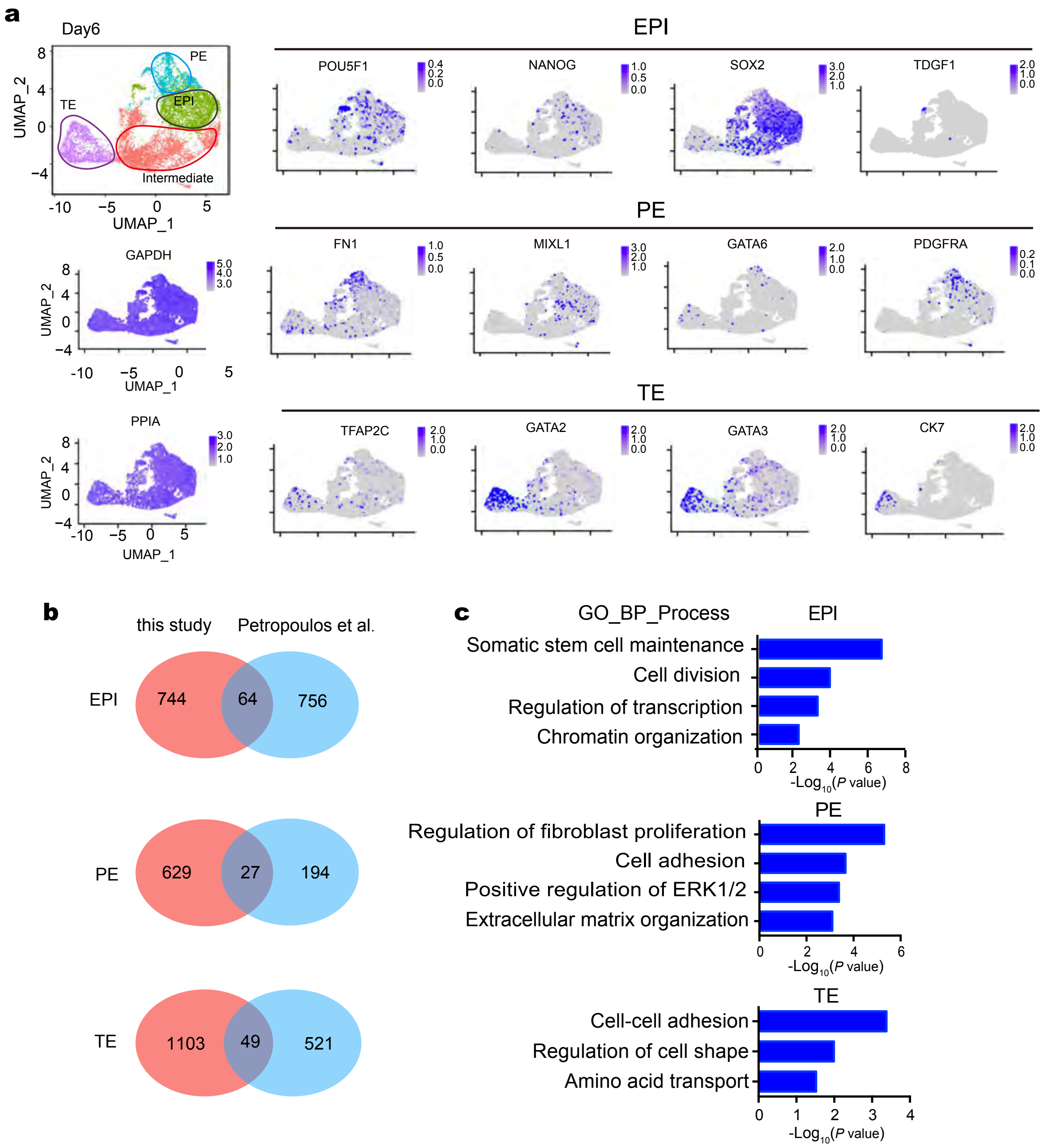


Extended Fig5

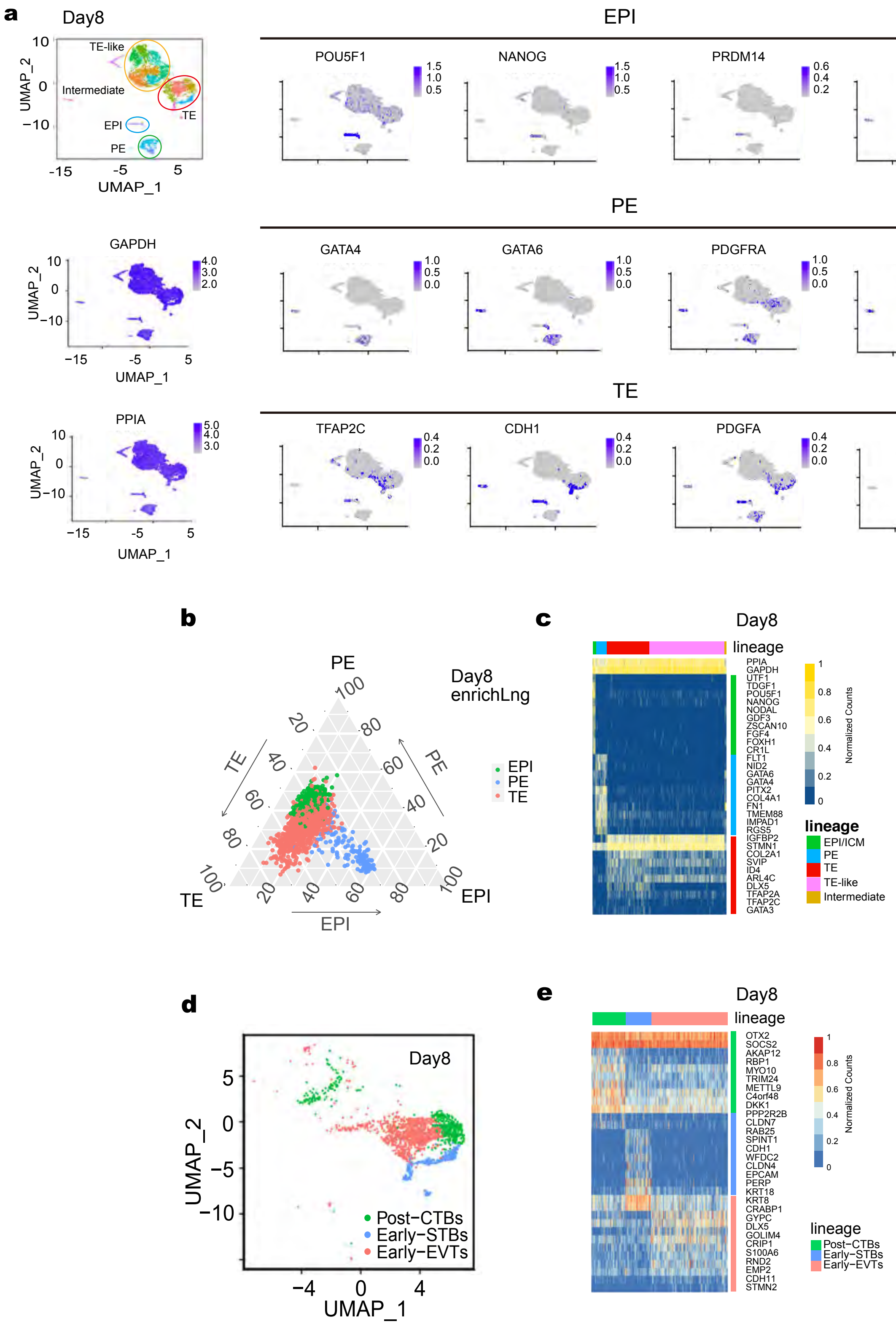


Extended Fig 6

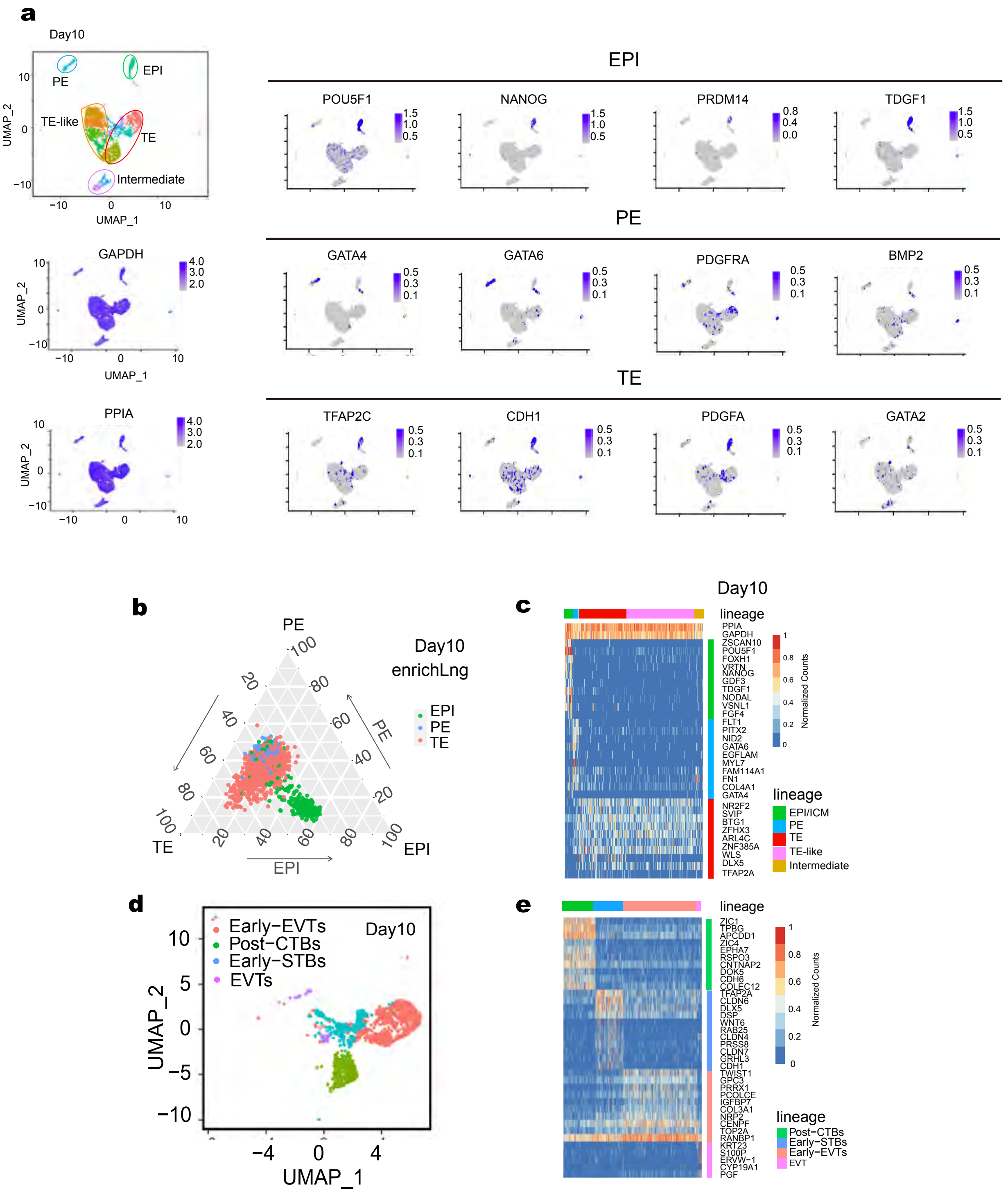


Extended Fig 7
